# Higher *in vitro* mucin degradation, but no increased paracellular permeability by faecal water from Crohn’s disease patients

**DOI:** 10.1101/2022.08.26.505386

**Authors:** Heike E.F. Becker, Nader Kameli, Alice Rustichelli, Britt A.M. Heijnens, Frank Stassen, John Penders, Daisy M.A.E. Jonkers

## Abstract

**Background:** Crohn’s disease (CD) is a chronic inflammatory gastro-intestinal condition with variable disease course. Impaired barrier function and microbial dysbiosis are associated with disease onset and exacerbations. We hypothesized that perturbed microbial activity may contribute to the impaired barrier function in CD. Therefore, this study aimed to examine the impact of faecal bacterial products of active and remissive CD patients, and healthy controls (HC) on mucin degradation and epithelial barrier function *in vitro*.

**Methods:** Six HC and twelve CD patients were included. Disease activity was determined by endoscopy. Fecal water (FW) and bacterial membrane vesicles (MVs) from fresh fecal samples were applied on mucin agar to determine mucin degradation and on differentiated Caco-2 cell monolayers to assess transepithelial electrical resistance (TEER) and paracellular junction stability. Relative abundances of fecal bacterial genera, which may be associated mucin degradation, were evaluated using 16S rRNA gene amplicon sequencing.

**Results:** FW-induced mucin degradation was higher in CD samples as compared to HC (p<0.01), but was not linked to specific bacterial relative abundances. FW resulted in 78-87% decrease of TEER in three of the remissive (p<0.001) but not the active CD or HC samples. MVs did not induce mucin degradation or epithelial barrier disruption.

**Conclusion:** The higher mucin degradation capacity of CD-derived FW might indicate contributions of microbial products to CD pathophysiology and warrants further investigation. Moreover, the altered epithelial resistance in some individuals is not due to paracellular disruption.

**Key Messages:** *What is already known?* Intestinal microbial dysbiosis and mucosal barrier dysfunction are important contributors to Crohn’s disease aetiology and disease exacerbations.

*What is new here?* The faecal microbial secretome of Crohn’s disease patients has a higher mucin degradation capacity as compared to the secretome of healthy subjects.

*How can this study help patient care?* The increased mucin degradation based on the microbial secretome may be a new target for the development of complementary, microbiome-based therapy in Crohn’s disease.

**Summary:** Microbial dysbiosis and intestinal barrier dysfunction can impact Crohn’s disease course. This translational study found higher mucin degradation, but no epithelial barrier disruption, by the faecal microbial secretome of (active) Crohn’s disease patients, as compared to healthy controls.

## Introduction

A balanced intestinal microbiota provides the host with several beneficial functions, such as immune modulation and defence against pathogens. In addition, microbes have a high metabolic activity, secrete a variety of metabolites, and release microbial peptides, which can interact directly or indirectly with the intestinal epithelium. For instance, short-chain fatty acids (SCFAs) or bacterial toxins, can directly interact with the mucus and epithelial layer and influence the leakage of luminal content to the host tissue. As an indirect mechanism, the release of microbe-associated molecular patterns (MAMPs), for example, stimulates the immune system via pattern recognition receptors, such as toll-like receptors, and can impact the immune status and mucosal integrity^1–3^.

The first site of host-microbe interaction is the mucus layer. In the colon, this consists of a loose outer layer, a tight inner layer and a membrane-anchored glycocalyx. The most abundant molecules in the mucus layer are mucins, which are O-glycosylated proteins excreted constantly by goblet cells^4^. The mucus layer is a crucial interface between the host and the intestinal microbes, amongst others by its high mucus turn-over and the presence of neutralizing immunoglobulins, whereas mucin glycoproteins also serve as nutrients for mucin degrading bacteria^4^. In turn, bacteria stimulate mucin production via MAMPs and the release of SCFAs^1^. In a healthy gut, mucin production and degradation are at equilibrium, while in intestinal diseases, there may be an imbalance^4^.

The second layer of the intestinal lining comprises the epithelial layer, which consists of multiple cell types and is apically sealed by tight and adherens junctions in the paracellular space. These limit the paracellular transfer of molecules from the intestinal lumen into the tissue^5^. Bacterial toxins, such as *Bacteroides fragilis* toxin or *Vibrio cholerae* hemagglutinin/protease and zonula occludens toxin, can disrupt these protein complexes resulting in increased paracellular permeability and thereby induce inflammation^3,4,6,7^. Further evidence suggests that bacterial membrane vesicles (MVs), containing different bacteria-derived molecules, including toxins, may also be able to disrupt paracellular junctions, as shown for *Bacteroides fragilis* toxin^8,9^. In contrast, bacterial products of saccharolytic fermentation, for example the SCFAs butyrate, propionate, or acetate, have been shown to improve epithelial connectivity by increased expression of several tight junction proteins, such as Occludin, Zonula Occludens protein-1 (ZO-1) and members of the Claudin family^2,10–12^.

A perturbed host-microbe interaction at the site of the mucosal layer has been shown to contribute to the pathophysiology of different diseases^4,13^. One example is Crohn’s disease (CD), which is a chronic inflammatory gastro-intestinal disease with alternating phases of remission (quiescent disease) and exacerbations (active inflammation)^14,15^. Several studies found microbial dysbiosis in CD patients compared with healthy controls (HC) and in patients during exacerbations as compared to those in remission. Such a dysbiosis is likely associated with a different composition of released microbial products, the secretome, thereby potentially contributing to a disturbed host-microbe interaction^16–19^. In addition, mucosal barrier dysfunction has been reported in CD patients, especially during exacerbations, which is characterized, among others, by altered tight junction expression and structures, and mucus secretion and composition^4,20^.

Increasing the knowledge about interactions between microbiota-released compounds and the mucosal barrier in situations of disturbed homeostasis, as observed in CD, will contribute to a better understanding of disease pathophysiology and may provide complementary therapeutic or preventive targets to improve disease outcome. Therefore, the aim of the present study is to explore the impact of faecal microbial products on the intestinal barrier *in vitro*, comparing the secretome of HC, active CD patients, and remissive CD patients. To this end, faecal water (FW) and faeces-derived MVs were examined for their ability to degrade mucin and disrupt the epithelial barrier.

The main findings of this study are a significantly higher capacity of mucin degradation by the faecal secretome of CD patients as compared to HC and inter-individual alterations in epithelial resistance.

## Methods

### Study population

Faecal samples were collected from six HC participating in the Maastricht IBS Cohort (MIBS) and twelve CD patients from the IBD South Limburg (IBD-SL) cohort^21^. The latter were grouped by their disease activity based on the Simple Endoscopic Score for Crohn’s Disease (SES-CD) with 0-2 points as remissive disease and ≥3 points as a current exacerbation^22^. Patients with prior colectomy, a current stoma or cancer were not eligible for this study. Faecal samples were collected within one week prior to bowel cleansing and endoscopy scheduled for clinical reasons.

### Faecal water isolation and dry weight

For maximum preservation of bacterial products, faecal samples were kept at 4°C and processed within 6 hours after collection. FW isolation was conducted following the protocol of Marchesi *et al*. with minor changes^23^. In brief, samples were homogenized and ∼15g was diluted 1:2 (w/v) in cold phosphate-buffered saline (PBS; Gibco, ref.:10010-031), vortexed for 1 minute and centrifuged at 4000x*g* for 15 minutes at 4°C instead of 3000x*g*. The supernatant was filtered through a 113V cellulose filter (cat no.:1213-150, Whatman, Germany) and subsequently centrifuged at 15000x*g* for 15 minutes at 4°C instead of 16000x*g*. The supernatant was sterile-filtered using 0.2µm pore size syringe filters with low protein binding (Sartorius, ref.:16534-5). The obtained FW was stored at -80°C until further analysis.

From frozen faecal samples, approximately 0.5g faeces were weighted and subsequently vacuum dried at 60°C for 5 hours in duplicate after which dry weight was determined.

### Membrane vesicle Isolation

For MV isolation, 1.5g defrosted faecal samples were diluted in 50ml cold PBS and centrifuged thrice at 4000xg for 15 minutes at 4°C. The obtained supernatant was filtered successively using syringe filters of 5µm (Pall, ref.:4650), 1.2µm (Pall, ref.:4656), 0.45µm (Pall, ref.:4653) and 0.2µm (Sartorius, ref.:16534-K), and subsequently concentrated to 250µl using Ultra Centrifugal Filters of 100kDa (Amicon Ultra-15, Merck Millipore, ref.:UFC9100) by repeated centrifugation at 4000xg for 15-60 minutes at 4°C. MV-containing fractions of size exclusion chromatography (SEC; sepharose CL-2B, ref.:GE healthcare, Little Chalfont, UK) were pooled and concentrations were measured by tunable resistive pulse sensing using qNano Gold (Izon Science Ltd., Oxford, UK) and Izon Control Suite software v3.2, as described by Benedikter *et al*.^24^

### Mucin degradation

To detect the capacity of mucin degradation by bacterial products of different faecal samples, we slightly adapted a previously published mucin degradation assay^25^. In short, 10µl FW or 10µl 10^8^ MVs were applied on 1.6% agar (ref.:LP0011, Oxoid) with 5g/l yeast extract, 5mg/l hemin and 37g/l brain heart infusion broth (ref.:53286, Millipore; BHI-YH), containing 0.5% partially purified porcine stomach mucin (ref:M1778, Sigma-Aldrich) and incubated anaerobically for 17 hours at 37°C. Mucin agars were then stained with Coomassie Brilliant Blue (Sigma-Aldrich) for 30 minutes and decolorized with 7% acetic acid and 7% methanol. Unfiltered faecal supernatant was used as positive control and sterile PBS as negative control. The amount of mucin degradation was determined by comparing the size of the translucent halo zone between the different FW or MV samples. The following scoring system was applied: 1=no halo zone, 2=halo zone smaller than halo zone of positive control, 3=halo zone as large as halo zone of positive control, 4=halo zone larger as halo zone from positive control. Each sample was tested in two independent experiments.

### Epithelial barrier evaluation

To investigate the effect of FW and MVs on intestinal epithelial permeability, Caco-2 cell monolayers (passages 47-57, RRID:CVCL_0025) were cultured in Millicell Hanging Cell Culture Inserts as well-established *in vitro* model for intestinal epithelial barrier function^26^. An initial density of 100,000 Caco-2 cells/insert was cultured for 14 to 21 days in Caco-2 culture medium (Dulbecco’s modified eagle medium ((DMEM; Sigma, ref.:D6429) supplemented with 1% (v/v) heat-inactivated foetal bovine serum (Gibco, ref.:10500), 0.1% (v/v) non-essential amino acids (Gibco, ref.:11140050) and 0.1% (v/v) antibiotic-antimycotic (Gibco, ref.:15240-062)) at 37°C and 5% CO_2_. Monolayers were exposed luminally to Caco-2 culture medium or 50% (v/v) FW diluted in Caco-2 culture medium. For MV evaluation, monolayers were exposed to Caco-2 culture medium with 50% PBS (v/v) with or without 10^8^ particles/ml MVs, diluted in Caco-2 culture medium. The transepithelial electrical resistance (TEER) was evaluated using the EVOM2 Epithelial Volt/Ohm Meter (World Precision Instruments, Sarasota, USA) of mature monolayers (TEER>600 Ω*cm^2^) prior to and after 24 hours of incubation. To further evaluate paracellular permeability, medium and culture conditions were replaced by 1mg/ml fluorescein isothiocyanate-labelled dextran of 4kDa (FITC-d4; Sigma) at the luminal site and PBS at the basal site. After 1 hour of incubation at same conditions, fluorescent intensity was measured at 485nm excitation and 530nm emission using a Spectramax M2 spectrophotometer (Molecular Devices). Each sample was tested for TEER and FITC-d4 flux in two independent experiments with three replicates. TEER results were expressed as percentage of initial TEER value and FITC-d4 permeation data as percentage of maximum permeation, which set by culture inserts without cell monolayers.

### Epithelial junction evaluation

To visualize localization of relevant tight and adherens junction proteins, Caco-2 monolayers were fixed with 4% (w/v) paraformaldehyde for subsequent immunofluorescent staining as described previously^27^. The procedure was conducted inside the transwell inserts and adjusted by initial permeabilization with 0.5% (v/v) Triton X-100 for 30 minutes and using PBS instead of Hank’s balanced salt solution for washing. Monolayers were incubated overnight with FITC-conjugated Zonulin-1 antibodies (1:200 dilution, ref.:339111, Invitrogen, RRID:AB_2533148) or anti-E-cadherin antibody (1:200 dilution, ref.:ab1416, Abcam, RRID:AB_300946). The latter was followed by three times washing with PBS for 5 min and incubation with goat anti-mouse IgG Alexa Flour 488 (1:200 dilution, ref.:A11029, Invitrogen, RRID:AB_138404) for 1 hour. Confocal images were taken with a Leica TCS SPE confocal microscope (Leica Microsystems GmbH, Mannheim, Germany) using LAS-AF software. Images were analysed qualitatively based on the morphology of the protein networks using ImageJ software. An intact junctional network was defined as cells being surrounded by a closed fluorescent lining and the morphology was comparable to the control condition.

### Faecal microbiota profiling

To identify associations between bacterial taxa (*i*.*e*. sulphate reducing and mucin degrading bacterial genera) and the observed mucin degradation, we profiled the faecal microbiota using 16S rRNA V4 gene region amplicon sequencing.

Faecal metagenomic DNA was isolated as published earlier^28^. Amplicon library preparation and sequencing was conducted as described previously^29^. Briefly, the PCR mix consisted of primers 515F and 806R (10pmol each), 5µl Accuprime buffer II, 0.2µl Accuprime Hifi polymerase (Thermo Fisher Scientific, Waltham, USA), 2µl DNA, and 32.8µl PCR-grade water. Upon amplification (3 minutes at 95°C, followed by 24 cycles of 30 seconds at 94°C, 45 seconds at 50°C, and 1 minutes at 72°C and a final step of 10 minutes at 72°C), amplicons were purified using AMPure beads (Agencourt, Massachusetts, USA) and quantified using Quanti-iT PicoGreen dsDNA kit (Invitrogen, New York, USA). Amplicons were then mixed in equimolar concentrations to reach ∼1ng/µl DNA and sequenced on an Illumina MiSeq using the V3 reagent kit (2×250 cycles).

The online integrated Microbial Next Generation Sequencing (IMNGS) platform with UPARSE-based analytical pipeline^30,31^ was used for demultiplexing (demultiplexer_v3.pl; unpublished Perl script), length and quality filtering, pairing of reads, and definition of Operational Taxonomic Units (OTUs) with 97% sequence similarity using USEARCH 8.1 with IMNGS default settings^32^. Chimera filtering was conducted using UCHIME (with RDP set 15 as reference database)^33^. Taxonomic classification was performed by RDP classifier version 2.11 and training set 15^34^. MUSCLE was used for sequence alignment and treeing by FastTree^35,36^. To exclude potential DNA contamination, negative extraction and amplification controls (PCR grade water) were compared to the lowest abundant microbial sample.

### Statistical analyses

Differences in mucin degradation between HC and CD samples were analysed using Mann Whitney-U test and between HC, remissive CD (R-CD), and active CD (A-CD) with Kruskal Wallis with Dunn’s post hoc test. Alterations in TEER and FITC permeation were analysed using Kruskal Wallis test when comparing groups and using ANOVA and Tukey’s post-hoc test when comparing individual samples. Correlations between mucin degradation and TEER alterations and faecal dry weight were tested using Spearman’s correlations. Differences in relative abundances among bacterial genera were analysed using Mann-Whitney test. Analyses were computed with GraphPad Prism 5 software and differences were considered statistically significant, when *p*<0.05.

### Ethical considerations

The study protocol was approved by the Medical Ethics Committee of the Maastricht University Medical Centre+ (MIBS: NL24160.068.08; IBD-SL: NL31636.068.10) and registered on www.clinicaltrials.gov (NCT00775060, resp. NCT02130349). All participants gave written informed consent prior to participation. The study was conducted in accordance with the revised Declaration of Helsinki and in compliance with good clinical practice.

## Results

### Patient population

Baseline characteristics of CD and control subjects (table 1) mainly differ with regard to medication use in CD patients. In addition, one HC used 200mg Ibuprofen at day two and four before sampling. Furthermore, one participant of each group followed a specific diet, which was pescatarian in one subject (HC) and low fat and reduced amounts of cabbage in the other (CD).

**Table 1:** Baseline characteristics of study participants.

|  | **Healthy control (n=6)** | **Remissive CD (n=6)** | **Active CD (n=6)** |
| --- | --- | --- | --- |
| **Gender (F/M)** | 5/1 | 5/1 | 3/3 |
| **Age (median (range))** | 41.5 (25-50) | 49.5 (38-67) | 47.5 (25-63) |
| **BMI (median (range))** | 22.7 (21.5-35.5) | 27.5 (23.4-34.1) | 26.0 (21.6-30.0) |
| **SES-CD (median (range))** | N/A | 0 (0) | 8.5 (3-21) |
| **HBI (median (range))** | N/A | 2.0 (0-12) | 4.0 (0-8) |
| **BSS (mean (range))** | 5 (3-5) | 4.8 (3-6) | 6.0 (5-6) |
| **Montreal Classification** |  |  |  |
| A1/A2/A3 | N/A | 0/4/2 | 1/3/2 |
| L1/L2/L3/L4 | N/A | 2/1/3/1 | 3/1/2/1 |
| B1/B2/B3/p | N/A | 5/0/1/1 | 4/1/2/2 |
| **Gastrointestinal surgery** | 0 | 2 | 3 |
| **IBD medication** |  |  |  |
| Mesalazine (n) | N/A | 1 | 0 |
| Budesonide (n) | N/A | 0 | 1 |
| Methotrexate (n) | N/A | 1 | 0 |
| Azathioprine (n) | N/A | 2 | 0 |
| Biologicals (n) | N/A | 1 | 5 |
| **Antibiotic use in past 3 months (n)** | 0 | 1 (claritromycin) | 2 (tobramycin, ciprofloxacin) |
| **PPI use (n)** | 0 | 1 | 1 |
| **Medication change** in last week<br>(n) | 1 | 1 | 0 |
| **Probiotic use** in last week (n) | 0 | 0 | 0 |
| **Alternative diet (n)** | 1 | 1 | 1 |
| **Current smoker (n)** | 0 | 1 | 2 |
| **Current pregnancy (n)** | 0 | 0 | 0 |

L1 = Terminal ileum; L2 = Colon; L3 = Ileocolon; A1 < 16 years, A2 = 17-40 years, A3 > 40 years at diagnosis; B1 = without structure formation, non-penetrating; B2 = Stricturing; B3 = Penetrating; BMI = body mass index; SES-CD = Simple endoscopic score of Crohn's disease; HBI = Harvey Bredshaw Index; BSS = Bristol Stool Scale PPI = proton pump inhibitor

### Faecal water and MV-induced mucin degradation

To detect differences in mucin degradation of faecal bacterial products, mucin containing BHI-YH agar was exposed to FW. After 17 hours of incubation, halo zones were observed for the positive control as well as for several FW samples with interindividual differences in diameter between samples, indicating mucin degradation (figure 1A; supplementary figure 1). The diameters were larger in A-CD (median score=3.25; IQR=3.00-3.88; *p*=0.022) compared to HC FW samples (median score=2; IQR=1.25-2.00), but were comparable to R-CD FW samples (median score=3.5; IQR=2.75-3.88; figure 1B). When comparing HC and CD, the diameters were significantly larger in CD FW samples (median score=3.50; IQR=2.88-4.00; *p*=0.007; figure 1C), indicating a higher mucin degradation capacity by CD-derived samples when compared to HC.

**Figure 1:**
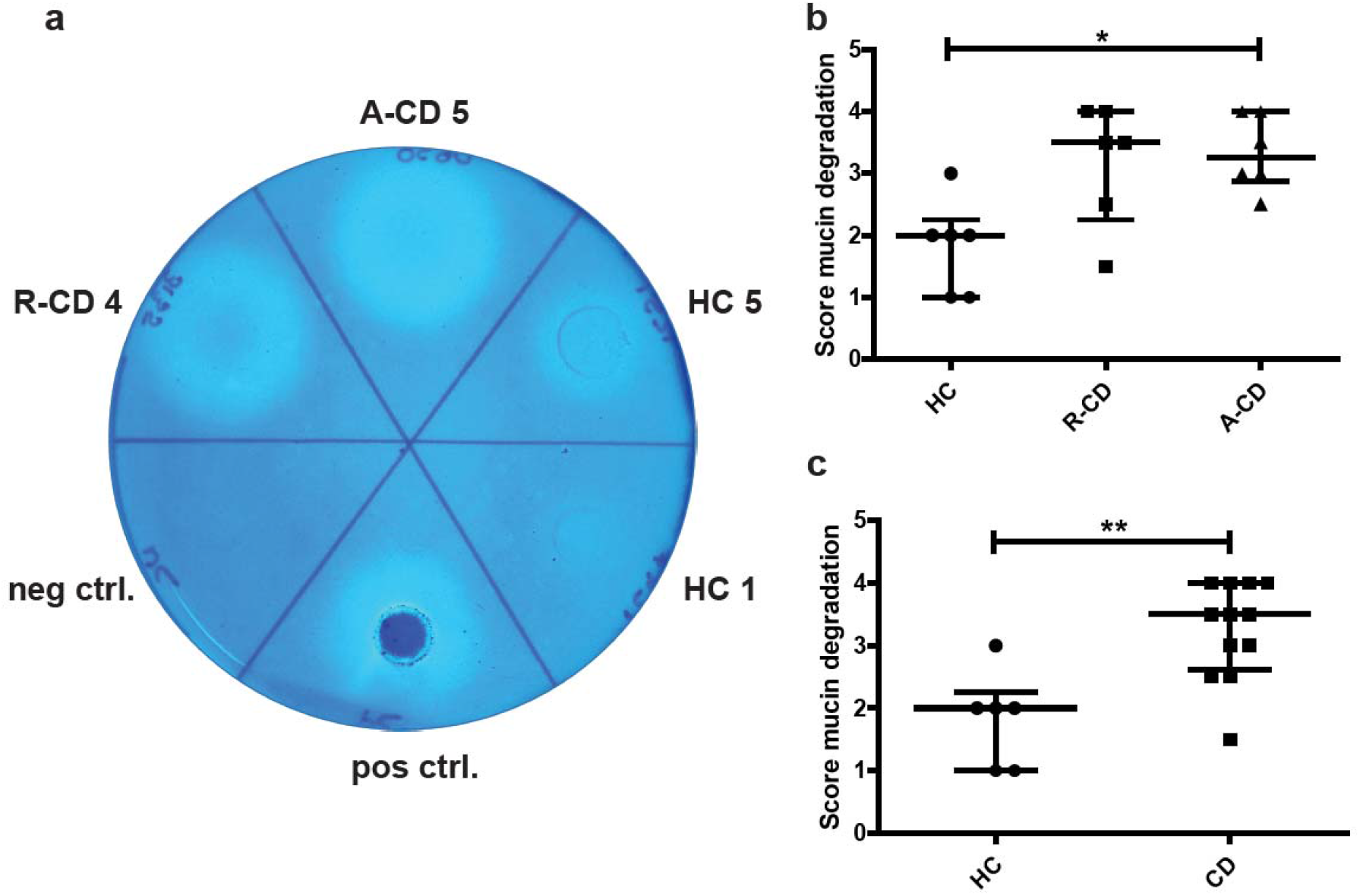
Fecal water of CD patients has a higher mucin degradation capacity compared with healthy subjects. (a) Selection of samples showing mucin degradation as indicated by the transparent halo zones. (b) Median scores of mucin degradation differed significantly comparing HC, R-CD and A-CD (*p* = 0.022; Kruskall-Wallis test). (c) Median mucin degradation scores were higher for FW of CD patients, compared to HC (*p* = 0.007; Mann-Whitney test). Each data point represents the average score of two independent experiments. Data are expressed as medians with IQR. * = *p* < 0.05, ** = *p* < 0.01; A-CD = active Crohn’s disease, R-CD = remissive Crohn’s disease, HC = healthy control

Concentrations of MVs ranged from 2.39×10^9^ to 2.78×10^9^ particles/ml and were diluted to a final concentration of 10^8^ particles/ml for each sample (supplementary figure 2). MVs did not induce mucin degradation as none of the MV samples resulted in detectable halo zones.

### Faecal water and MV-induced barrier alterations

To investigate the impact of faecal bacterial metabolites on intestinal epithelial barrier function, FW (50% v/v) was applied on Caco-2 cell monolayers in a transwell system. After 24 h incubation, no significant alterations in TEER were detected between HC, active, and remissive CD patients (figure 2A). However, the group of remissive CD patients showed a high standard deviation due to patient-specific variation of individual samples; three remissive patient samples showed no effect, whereas samples from another three remissive patients showed an 82.6-87.4% decrease in TEER values compared to negative control (*p*<0.001, figure 2B). No evident differences in demographical, including age and sex, or patient characteristics, including HBI and medication use, could provide explanations for the detected TEER decrease. For none of the investigated samples, alterations in FITC-d4 permeation could be detected (figure 2C, D), which indicates that the epithelial barrier remained intact upon FW exposure.

**Figure 2:**
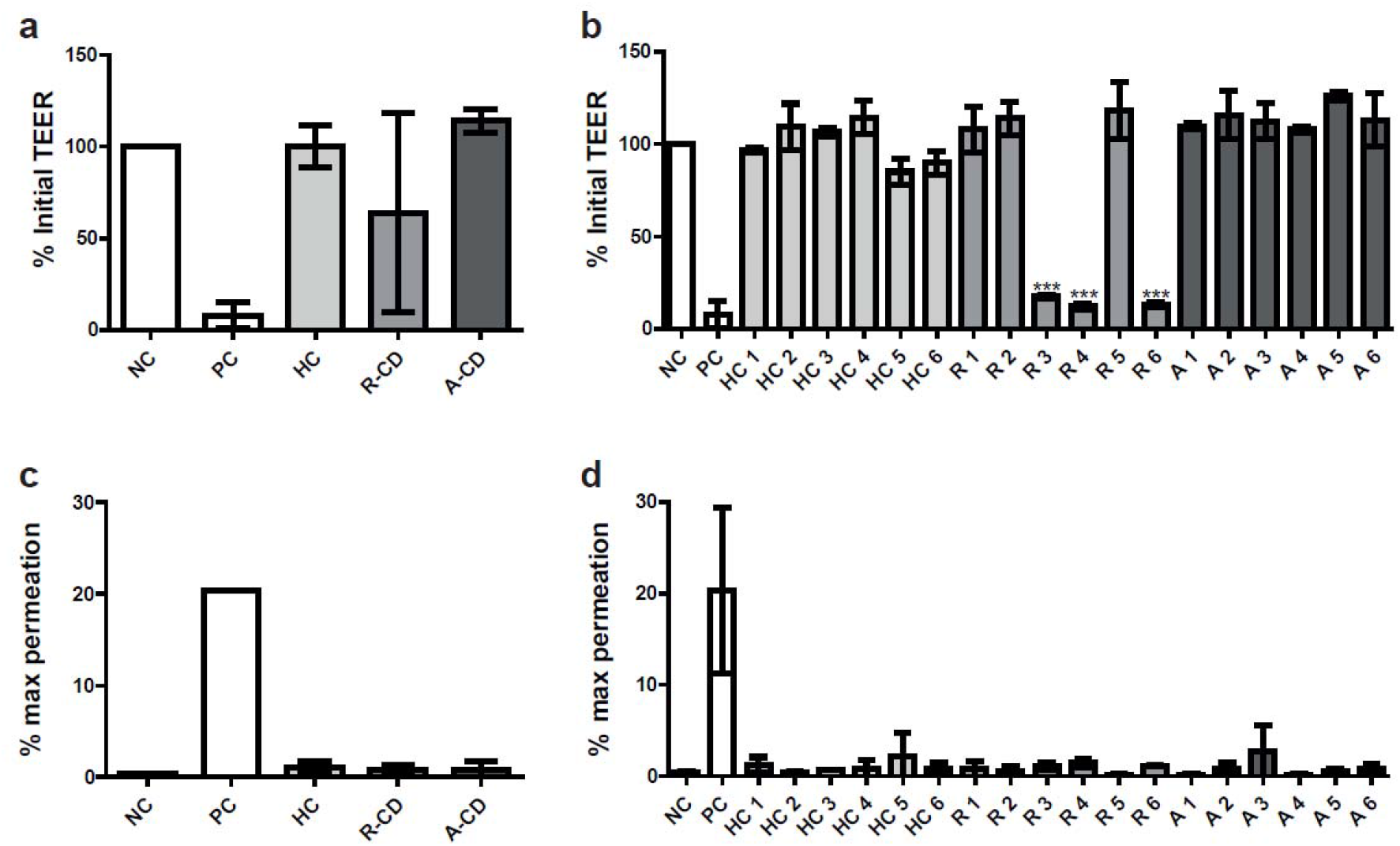
Luminally applied fecal water alters TEER on individual basis. (a) TEER values did not alter significantly by exposure to FW (50 % v/v) after 24 hours. (b) TEER is 82∙6-87∙4 % decreased in response to FW of three R-CD patients (*p* < 0.001). (c, d) No significant alterations were detected by FITC-d4 permeation on group (c) or individual level (d). Data are expressed as means ± SD. Group analyses were conducted using Kruskal-Wallis and analyses for individual sample using One-way ANOVA and Tukey’s post hoc test without the positive control. *** = *p* < 0.001; NC = negative control; PC = positive control; HC = healthy control; A/A-CD = active CD; R/R-CD = remissive CD

Isolated MVs did not affect TEER nor FITC-d4 permeation levels (figure 3).

**Figure 3:**
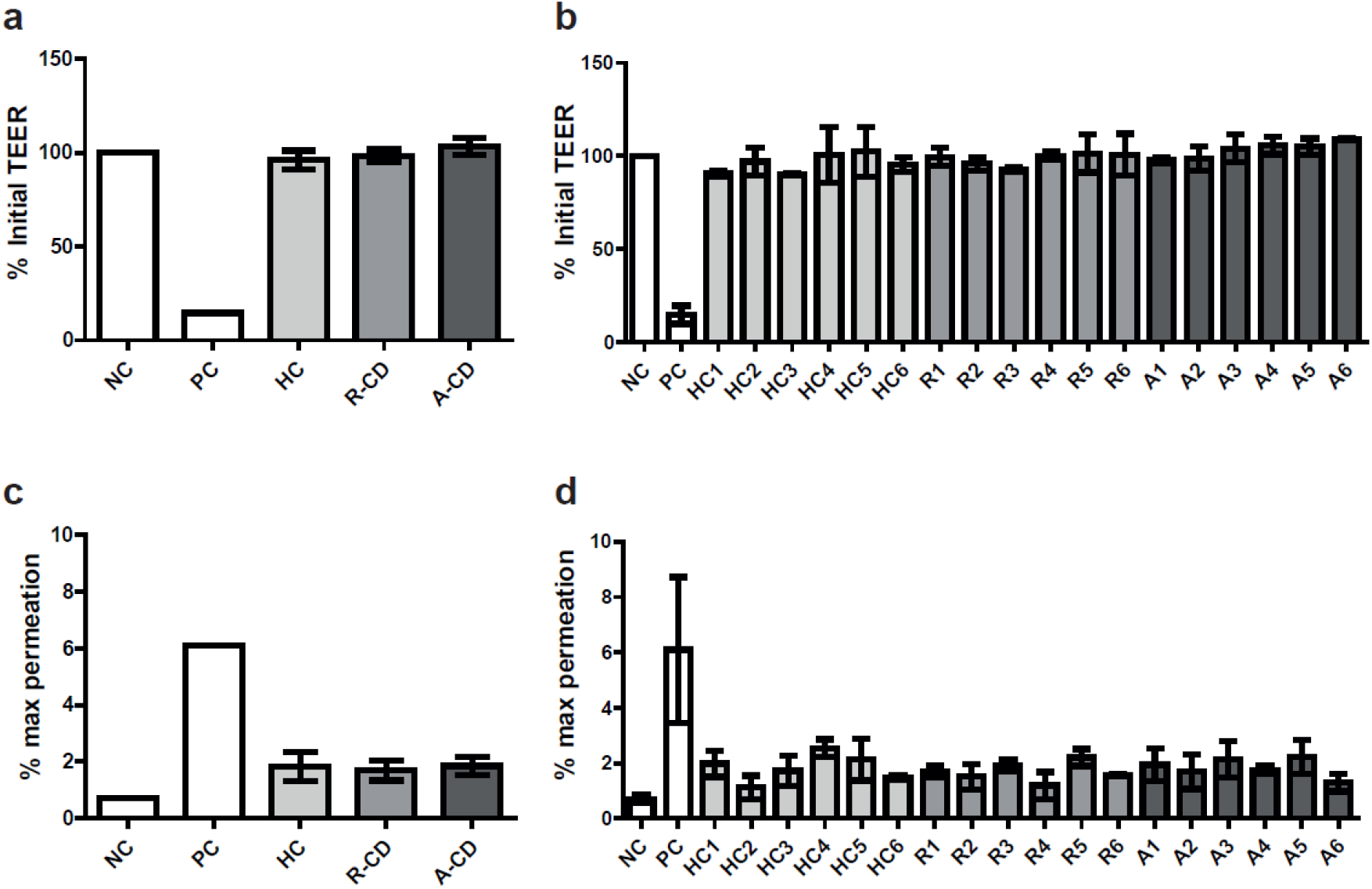
TEER remains unaltered in response to luminal application of MVs. (a, b) TEER is comparable in response to luminally applied 10⁸ MVs/ml on group (a) and individual (b) level after 24 hours. (c, d) FITC permeation is comparable on group (c) and individual (d) level with the same conditions. Data expressed as means ± SD. Group analyses were conducted using Kruskal-Wallis and analyses for individual sample using One-way ANOVA and Tukey’s post hoc test without the positive control. NC = negative control; PC = positive control; HC = healthy control; A/A-CD = active CD; R/R-CD = remissive CD

### The impact of faecal water on junctional protein localization

To further exclude an impact of FW on the paracellular junctional complex, the following six samples were selected to investigate the morphology of the tight junction protein Zonulin-1 and the adherens junction protein E-cadherin by immunofluorescence: three samples with a significant TEER decrease (R-CD 3, 4, 6), two samples with a marginal increase (A-CD 2, 5), and one sample without alteration in TEER (A6). Both proteins showed continuous junctional networks in all samples indicating intact tight and adherens junctions (figure 4).

**Figure 4:**
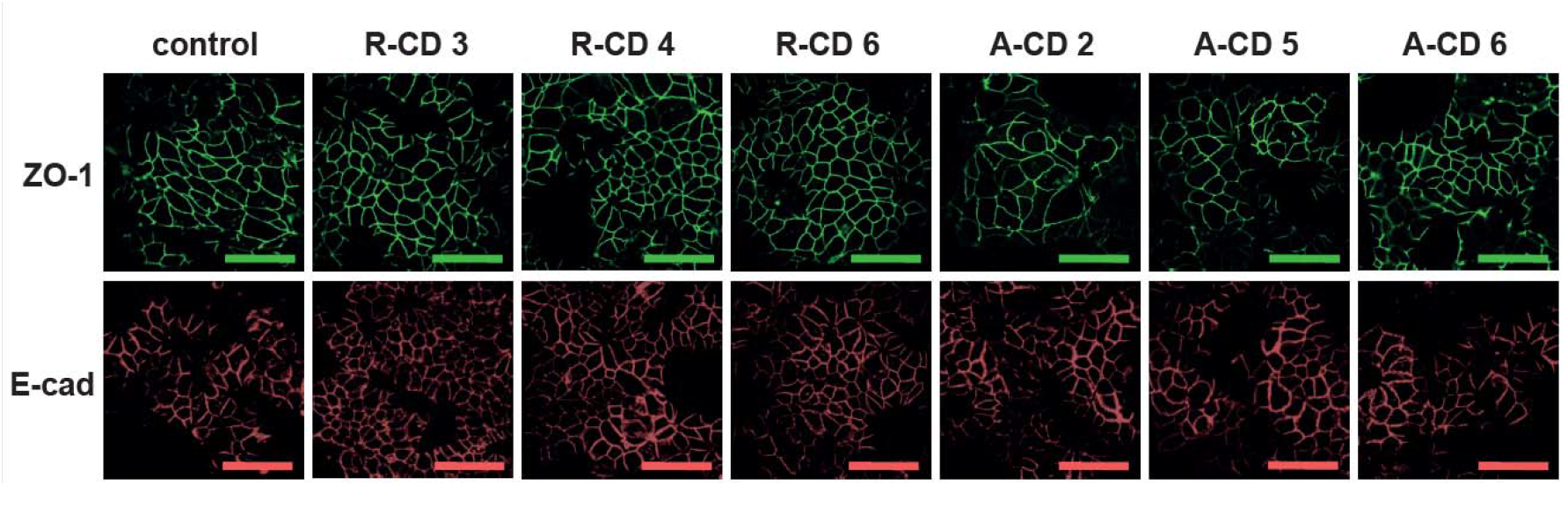
Representative tight and adherence junctions remain unaltered in Caco-2 monolayers in response to fecal water of CD patients. All samples show comparable characteristic morphological patterns of the immunofluorescently labelled tight junction protein ZO-1 and the adherens junction protein E-cadherin on confocal microscopic images after 24 hours incubation with FW (50 % v/v). This indicates intact junctional connections in all investigated conditions, including R-CD 3, 4, and 6, which had decreased TEER values. Scale bar = 40 µm; A-CD = active CD; R-CD = remissive CD

### The impact of faecal dry weight on mucin degradation and TEER

To exclude any impact of faecal consistency on experimental outcomes, the correlation of faecal dry weight with mucin degradation scores and TEER was investigated. Using Spearman’s correlation, no correlation with the percentage of faecal dry weight was found neither for mucin degradation (ρ=-0.444; figure 5A) nor for alterations in TEER (ρ=-0.259; figure 5B).

**Figure 5:**
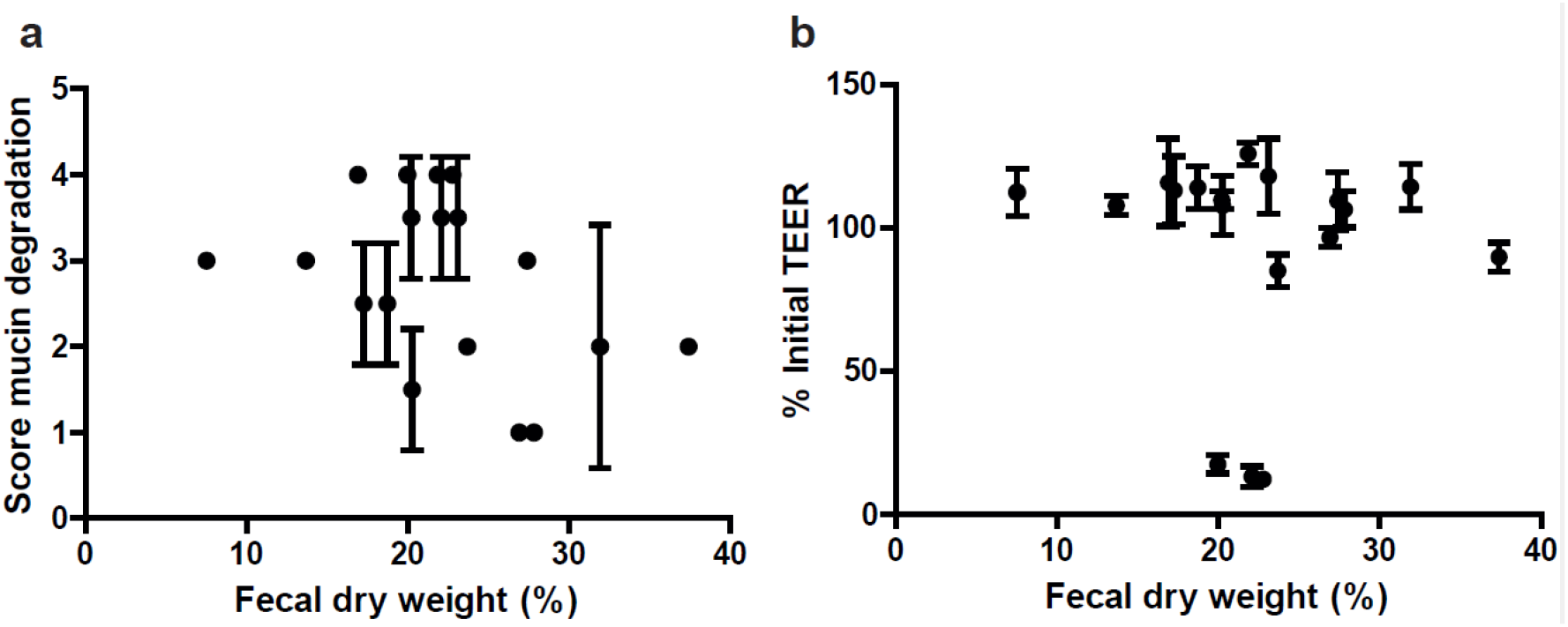
The dry matter does not influence the effects of fecal water on mucin degradation and TEER. (a) Fecal dry weight does not correlate with the scores of mucin degradation (ρ = -0.444) and (b) the percentage of initial TEER values (ρ = -0.259). Data expressed as means ± SD.

### Relative abundances of bacterial genera were comparable among groups

To identify potential associations between specific bacteria and mucin degradation capacity by FW of HC and CD subjects, relative abundances of faecal bacterial genera were analysed. However, no significant differences were detected comparing HC with CD, especially with regard to genera which contain known mucin degraders, such as *Akkermansia* and *Bacteroides*, as well as sulphate reducers, such as *Desulfovibrio* and *Bilophila* (figure 6). Additional information on the microbiota composition in the investigated samples can be found in the recent publication by Kameli et al. (2021) ^37^.

**Figure 6:**
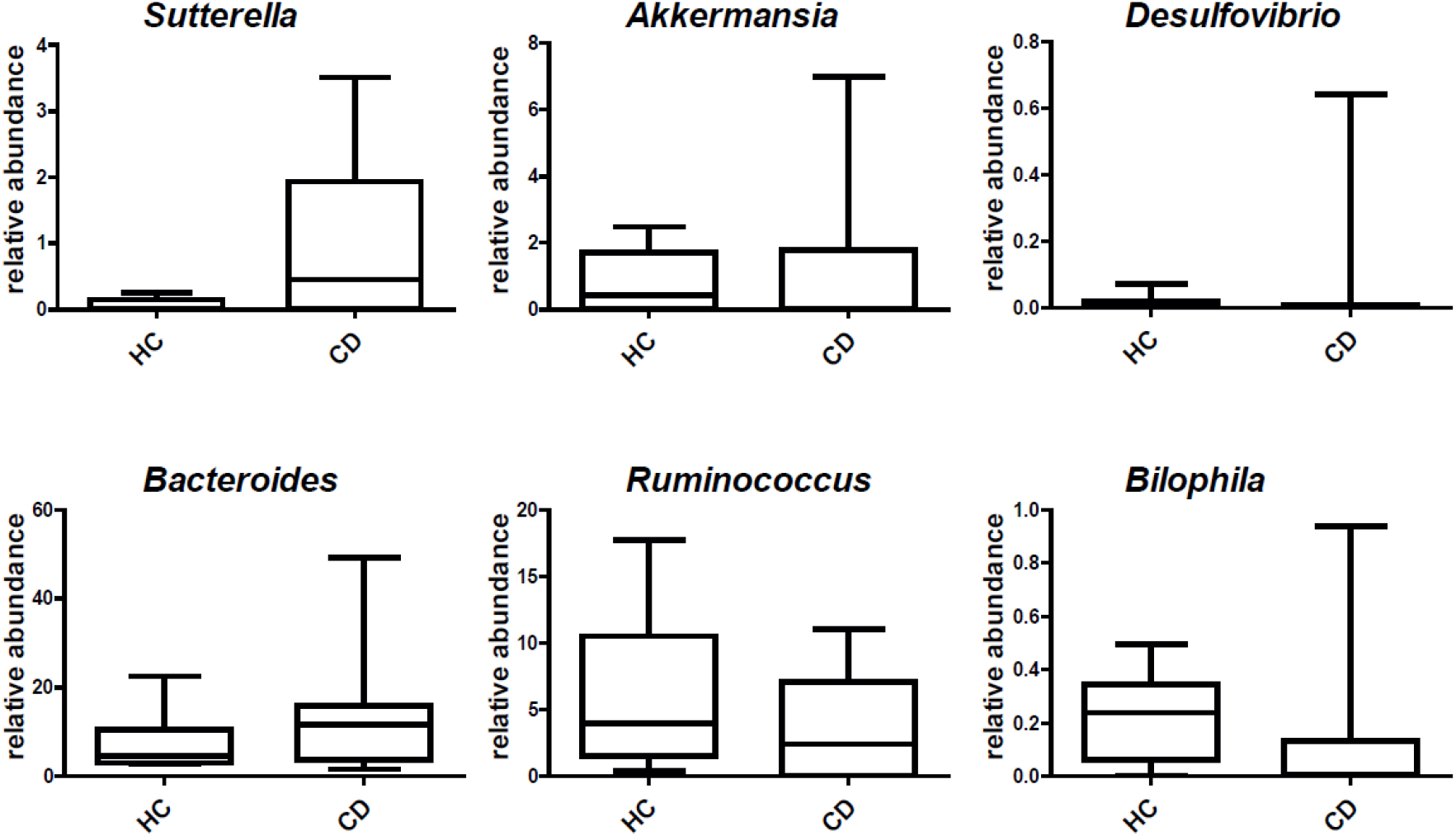
Bacterial genera do not differ between HC and CD. Bacterial genera did not differ significantly in relative abundances when comparing faecal DNA of HC and CD patients. Data of selected genera are presented as median + IQR.

## Discussion

Previous research highlighted intestinal dysbiosis and mucosal barrier dysfunction as important factors contributing to CD pathophysiology and disease course^13,14,16,17,20^. Based on the reciprocal interaction of intestinal microbes with the host mucosal barrier^1,4^, we hypothesized that dysbiotic alterations of the faecal microbial secretome could impair barrier function and thereby contribute to exacerbations in CD. In this study, both, the active and the remissive CD secretome increased mucin degradation when compared to the secretome of HC. Furthermore, we found stable paracellular junctions in epithelial monolayers exposed to the secretome of all subjects, but inter-individual differences on TEER.

We used FW to represent the colonic microbial secretome, which can interact with the host mucosal tissue. Previous research repeatedly found a perturbed microbiota composition in CD patients as compared to HC, which was characterized by a lower microbial diversity and richness, lower temporal stability, and several taxonomic differences, such as a decrease in Firmicutes and an increase in Proteobacteria^14,16–19^. This makes the composition of the microbial secretome in CD patients likely to be different as well^38^. Using FW, we observed more mucin degradation by active CD patients’ samples compared to HC. Although a previous study has found that bacterial MVs contribute to mucin glycan degradation, which facilitates further degradation by bacteria and serves as nutrient source, we did not detect any mucin degradation by MVs^39^.

For mucin degradation experiments, the current study used porcine gastric mucin. This type of mucin is similar to human MUC5AC, which can be found in the human stomach^40,41^. Although, this type of mucin is not identical to colonic MUC2, it has been used previously for representative *in vitro* studies^25,41,42^.

Since only a few bacterial taxa are able to degrade mucin^43^, the higher mucin degrading capacity by the CD patients’ secretome may originate from a higher abundance of mucin degraders and therefore higher amounts of secreted mucinases^43,44^. In addition, a higher abundance of sulphate reducing bacteria (SRB) was found previously in CD patients compared to HC^45^. SRBs use sulphate as energy source, generating hydrogen sulphide, which reacts with the mucin disulphide bonds and subsequently degrades the mucin network^46^. However, we did not detect differences in the relative abundance of specific faecal bacterial genera, neither among known mucin degraders, such as *Akkermansia* and *Bacteroides*, nor among SRBs, such as *Desulfovibrio* and *Bilophila*.

Therefore, it seems that taxonomic composition might not reflect the function of the secretome regarding mucin degradation. Instead, several factors can influence the microbial mucin degradation capacity. For instance, the mucin degrader *Bacteroides thetaiotaomicron* prefers dietary polysaccharides, but increases its mucin degradation capacity during low dietary polysaccharide availability^47^. This implies that the degradation capacity might be affected by differences in dietary habits and thus substrate availability. Furthermore, intestinal microbes collaborate in mucin degradation with few taxa being able to utilize the mucin proteins after sialic acid removal and oligosaccharide degradation by others. For instance, secreted α- and β-glucosidases by *Ruminococcus gnavus* and *Ruminococcus torques* can remove the terminal sugars, thereby facilitating mucin degradation^48^. This highlights the complex cross-feeding interactions between different members of the microbiome. Together, these factors indicate that the microbiota composition alone may not explain the observed difference in mucin degradation by the studied microbial secretome. The interaction of intestinal microbes and additional causing factors of mucin degradation in CD remain to be explored. Besides an altered microbiota composition, previous studies found an altered mucin structure in CD, which is likely due to altered levels of transcription and post-transcriptional mucin^4,44^. For instance, CD patients display a higher *MUC2* expression, reduced length of the mucin associated oligosaccharide chains, and increased sialylation^44^. Since bacterial mucin degraders depend on glycosylated mucins as direct nutrient source and other microbes use their secreted metabolites, it cannot be excluded that the observed microbial perturbations in previous studies are a consequence rather than a cause^4,43^. This potential niche effect might also explain why we did not detect a difference in mucin degradation capacity between active and remissive CD samples.

However, in case of a causal role of the microbiota in mucin degradation, this can facilitate a closer contact of microbes with the mucosal layer and stimulate epithelial damage and subsequent inflammation.

In addition to the mucus layer, the underlying epithelial layer, which is sealed by tight and adherens junction proteins, plays an important role in intestinal barrier function^5^. Investigating the direct effect of FW on epithelial integrity, we found no significant decrease in TEER values after exposure of Caco-2 monolayers to FW of CD patients and HC. However, three samples from remissive CD patients did lead to a significant decrease in TEER, which might rather be an individual than disease-specific effect. Interindividual differences were not conspicuously linked to faecal consistency or baseline characteristics, such as medication use, smoking, or BMI. As dietary intake was not analysed in the current study, we cannot exclude a potential impact of food particles on TEER or mucin degradation. The observed decreases in electrical resistance were not accompanied by an increased FITC-d4 flux nor pattern alterations of stained ZO-1 and E-cadherin, and thereby not indicative for increased paracellular permeability for small or large molecules^49^. Instead, the individual alterations in TEER might be due to differences in ion flux by faecal metabolite and ion contents^26,50^. Herein, ion channel forming Claudin proteins, such as Claudin-2, -4, 12, or -15, might be modulated with regard to expression, post-transcriptional modification, or localisation^51^. Potential consequences and relevance with regard to tissue fluid homeostasis and inflammation in CD remain to be investigated. Although we could not detect bacterial secretome-induced tight and adherens junction modification, intestinal bacteria can still exert a direct effect. This was previously shown for *Bifidobacterium longum* and *B. bifidum* strains^2^. Therefore, future studies should expand on the investigation of direct bacteria-epithelium interactions to better understand the functional effects of microbial dysbiosis in CD.

In the current study, we used a well-accepted Caco-2-based model to study intestinal barrier function, which is based on a colon cancer cell line^26^. Future examinations of current findings may translate our approach to a more physiological model, such as intestinal disease-specific non-cancer organoids^52^, which could be of additional relevance. Furthermore, our findings provide new leads to study additional indirect effects of the CD patient faecal secretome, such as immune-mediated effects applying co-culture models including relevant immune cells. In addition, larger CD patient cohorts may be screened, which further allows the examination of specific disease phenotypes in relation to faecal secretome function.

Taken together, we found a significant impact of the microbial secretome of CD patients on intestinal mucin degradation but not on epithelial permeability *in vitro*. It remains to be elucidated whether the higher mucin degrading capacity in CD samples is a contributor to barrier disruption or a niche-response to the CD-related altered mucus composition and whether the dysbiotic microbiota of CD patients has a direct functional effect on the epithelial barrier.

## Acknowledgments

H.B. was funded by a NUTRIM Graduate Programme Grant from Maastricht University.

