## Supplementary Figure 1 for "Higher *in vitro* mucin degradation, but no increased paracellular permeability by faecal water from Crohn’s disease patients"

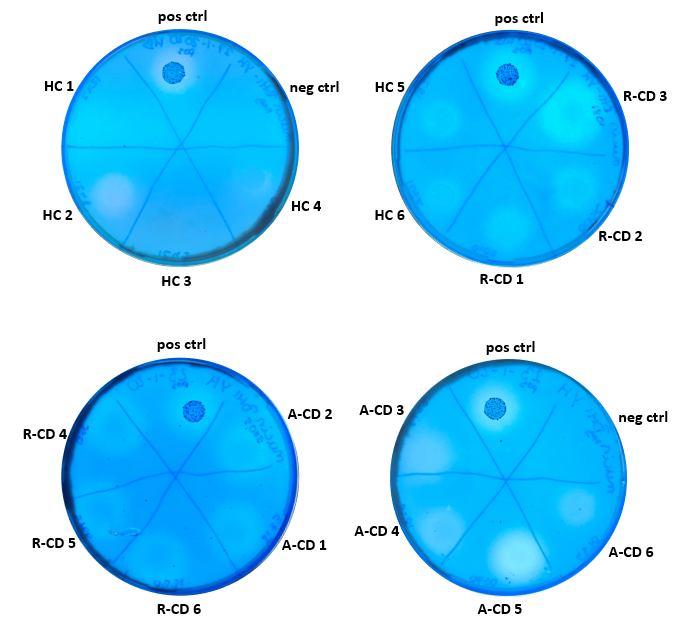

**Supplementary Figure 1: Mucin degradation by all samples.**

Mucin degradation varies as indicated by the transparent halo zones. A fresh, unfiltered faecal water sample was used as positive control (pos ctrl) and sterile PBS was used as negative control (neg ctrl).

HC = healthy control, R-CD = remissive Crohn’s disease, A-CD = active Crohn’s disease
