## Supplementary Figure 2 for "Higher *in vitro* mucin degradation, but no increased paracellular permeability by faecal water from Crohn’s disease patients"

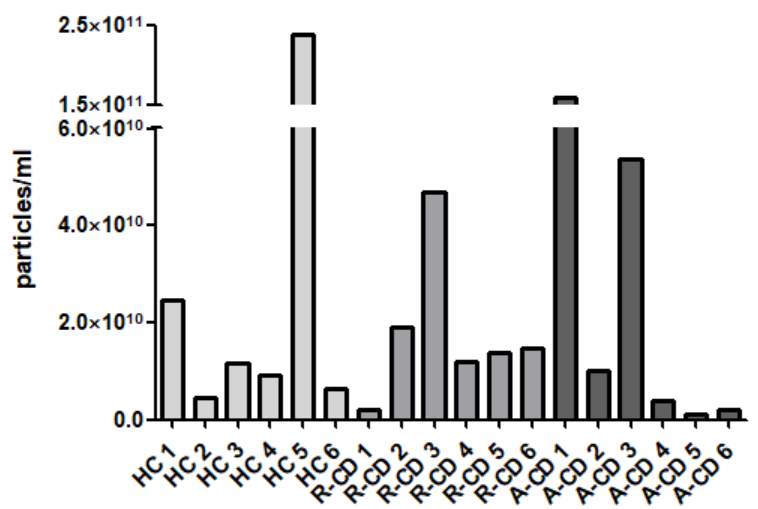

**Supplementary Figure 2: Detected concentrations of MVs**

Each sample was measured individually and showed successful isolation of bacterial MVs. 1 ml of isolated vesicles corresponds to 1.5g faecal sample.

HC = healthy control, R-CD = remissive Crohn’s disease, A-CD = active Crohn’s disease
